# Developmental staging and future research directions of the model marine tubeworm *Hydroides elegans*

**DOI:** 10.1101/2022.10.24.513551

**Authors:** Katherine T. Nesbit, Nicholas J. Shikuma

## Abstract

**Background:** The biofouling marine tube worm, *Hydroides elegans*, is an indirect developing polychaete with significance as a model organism for questions in developmental biology and the evolution of host-microbe interactions. However, a complete description of the life cycle from fertilization through sexual maturity remains scattered in the literature, and lacks standardization.

**Results:** Here we present a unified staging scheme detailing the major morphological changes that occur during the entire life cycle of the animal. These data represent a complete record of the life cycle, and serve as a foundation for connecting molecular changes with morphology.

**Conclusions:** The present descriptions and associated staging scheme are especially timely as this system gains traction within research communities. Characterizing the *Hydroides* life cycle is essential for investigating the molecular mechanisms that drive major developmental transitions, like metamorphosis, in response to bacteria.

## INTRODUCTION

The marine tube worm *Hydroides elegans* is an emerging model Spiralian for addressing questions in developmental biology and probing the molecular foundations of bacterial influences on development. *Hydroides* is a simple organismal model that is highly tractable in a laboratory setting. They release thousands of eggs and sperm at a time, and spawning is non-lethal. Fertilization occurs externally, and the embryos/larvae will rapidly develop in a simple dish of sea water. Metamorphosis of *Hydroides* larvae will occur at five days, and is catalyzed by contact with a heterogeneous bacterial biofilm. In the lab, metamorphosis can be recapitulated by contact with axenic biofilms composed of a single stimulatory bacterial strain [8, 10, 11].

*Hydroides* enables single animal-single microbe investigations, ultimately facilitating characterization of the pathways through which eukaryotic organisms sense and respond to an array of microbial products that induce major developmental transitions.

Despite its utility as a developmental model[12-17], characterizations of the growth and development of *Hydroides* remains fragmented across the existing literature[12, 14-18]. These works include partial descriptions of morphological and molecular features at the cleavage stages[14] and early embryogenesis[15], as well as formation of the feeding larval stages[16], segmentation[12], and emphasis on metamorphosis and juvenile life stages[8, 9, 17-19]. Yet, the community has not established a standardized staging scheme that spans the entirety of development. To this end, we provide here a complete staging of the life cycle of *Hydroides* (Figure 1) from fertilization through sexual maturation, as well as offer perspectives on how this model can be leveraged to advance our understanding of diverse and fundamental biological questions.

**Figure 1.**
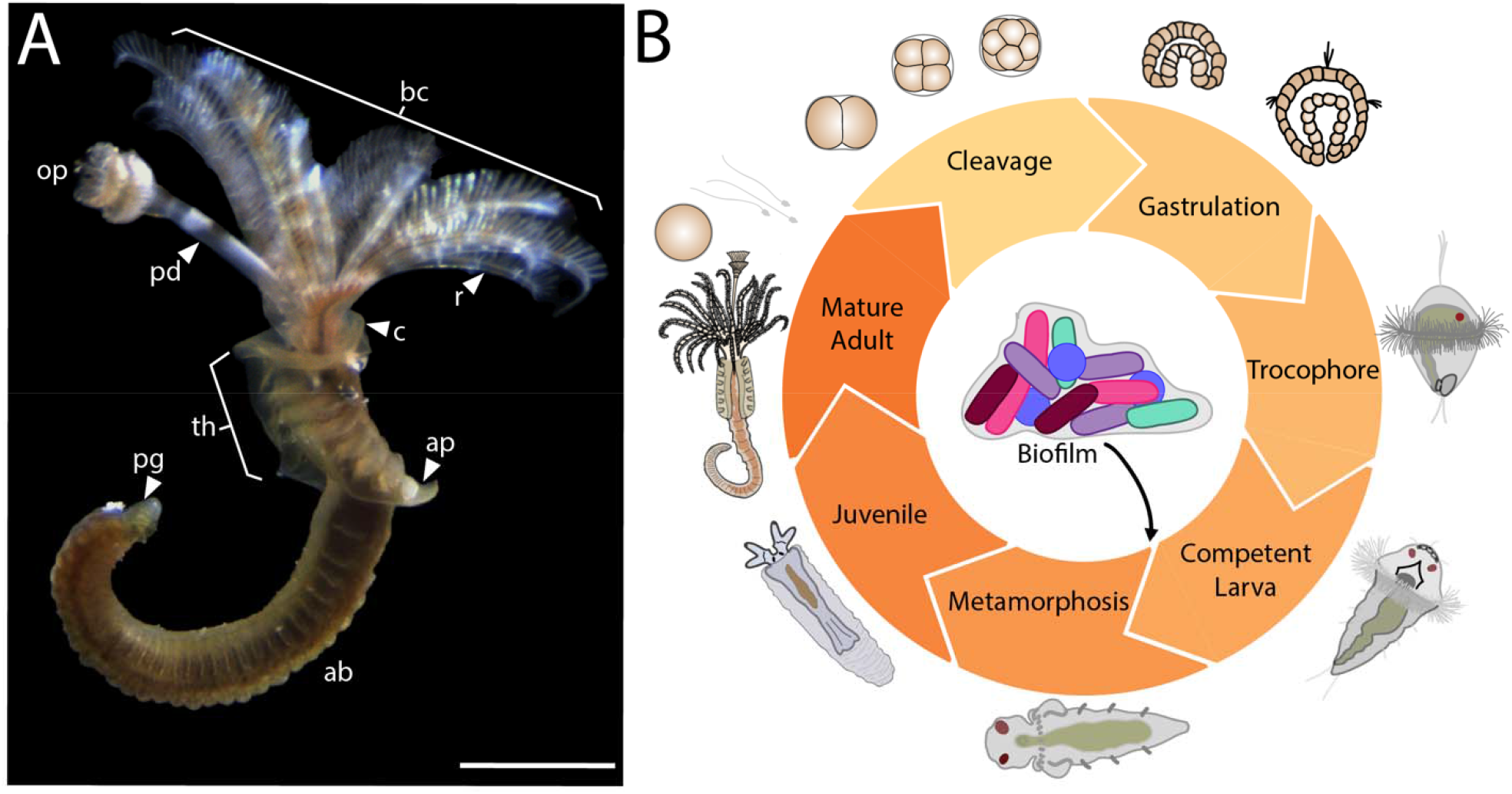
*Hydroides* morphology and life cycle. A) Sexually mature adult dissected out of the calcareous tube. Labelling: ab=abdomen, bc=branchial crown, c=collar, op=operculum, pd=peduncle, pg=pygidium, r=radiole, th=thorax. B) Illustration of the major life history stages in the biphasic life cycle of Hydroides elegans. Metamorphosis is induced by interaction with single- or multi-species biofilms. Scale = 1mm.

## EVOLUTION AND NATURAL HISTORY OF HYDROIDES

Annelids are a diverse and abundant group of organisms, and are part of a larger evolutionary clade known as the Spiralia[39]. Spiralians are of particular interest in evolution and development because of the vast diversity of bilateral body plans they exhibit throughout ontogeny, which arise from a highly conserved embryonic cleavage pattern. The Annelids represent an under-sampled and understudied source of Spiralian variation in the evolution of development, despite having been shown to exhibit several important developmental differences to molluscs[38, 40]. Existing annelid models with developmental staging schemes include the leech – *Helobdella*[26], the ragworm – *Platynereis*[27], the parchment worm - *Chaetopterus*[28], *Capitella*[12], and *Owenia*[29]. As a Spiralian developmental model, *Hydroides* can help us understand the evolution of innovation and conservation in development at the cellular and molecular level, as well as how this intersects with environmental input in development within and among major metazoan lineages.

Adult *Hydroides* (Figure 1A) are protandrous hermaphrodites, and are found in intertidal to sublittoral zones throughout the global tropical and subtropical oceans[41, 42]. They are part of a larger group of tube-building annelids in the family Serpulidae. These animals are filter feeders, extending branching appendages, collectively called the branchial crown, out from the opening of their tubes to collect particles from the water. In addition, they have a special funnel-shaped structure called an operculum that closes off the opening to the tube when the animal retracts inside. As one of the most prolific biomineralizing annelids, this species has garnered attention as a biofouling organism that deposits its calcareous tubes on ship hulls and other submerged surfaces in harbors and marinas[11, 42, 43]. They are resilient, tolerating variable temperatures (15-30°C) and salinities (15-37 PSU)[13], and able to colonize surfaces treated with different antifouling agents[44, 45]. However, despite being seen as “pests” in harbors and marinas, these animals are valuable research models. They, along with other members in the genus, have been utilized for biological research since the species was first described over 100 years ago[46], with increasing interest over the last few decades. This surge in interest in *Hydroides* centers around antifouling and the biphasic life history (Figure 1B) of this animal which utilizes microbial intervention to complete development[8].

## DEVELOPMENT

Unlike the gametes of other marine invertebrates, for example sea urchins, there is no obvious morphological changes to the eggs of *Hydroides* upon fertilization (Figure 2). The early cleavages of *Hydroides* (Figure 3) are equal-sized[14]. Unlike other members of the Spiralia and even other Annelids[14, 37, 38], the “D” quadrant of the embryo is not morphologically distinct from the other regions at the 4-cell stage (Figure 3B). The third cleavage, yielding an 8-cell embryo (Figure 3C), is the first division during which the spiral pattern becomes apparent. This division is a sinistral (left-handed) twist[14]. The cleavage spindle rotates, and as a result the daughter cells are born on the other side of the macromere, giving the appearance that the blastomeres on the animal side of the embryo have rotated counter-clockwise relative to the vegetal blastomeres[14]. The 8 cells of the embryo complete the next cleavage cycle concurrently, yielding the 16-cell stage (Figure 3D). This is in contrast to the blastomere divisions leading from 16- to 32-cell stage (Figure 3E), and the 32- to 64-cell stage (Figure 3F) transition which do not occur synchronously[14].

**Figure 2.**
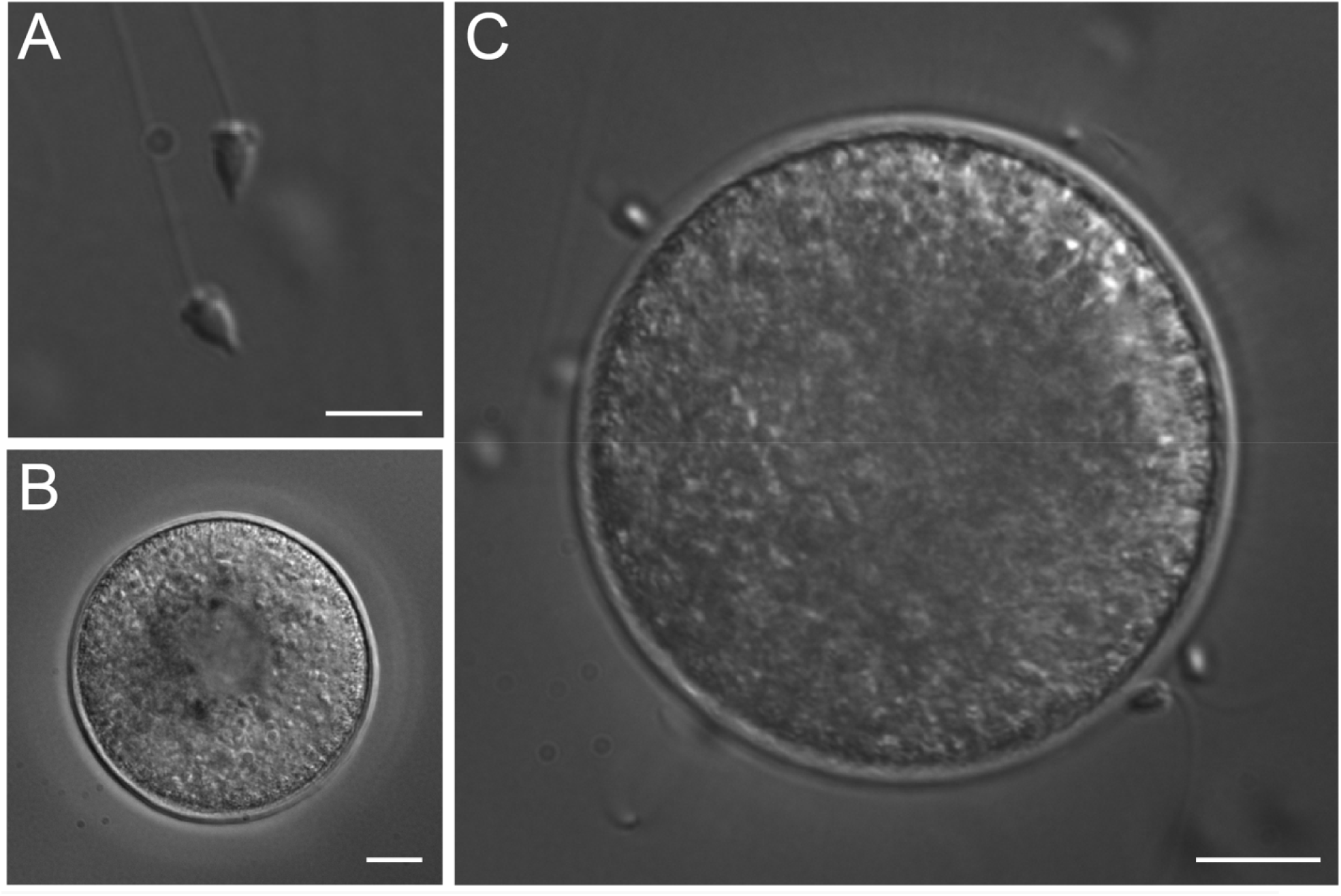
Gametes and zygote of *Hydroides elegans*. A) Sperm. Scale=5μm. B) Unfertilized egg. Scale=10μm. C) Zygote. No obvious morphological changes occur between the oocyte and the zygote. Scale=10μm.

**Figure 3.**
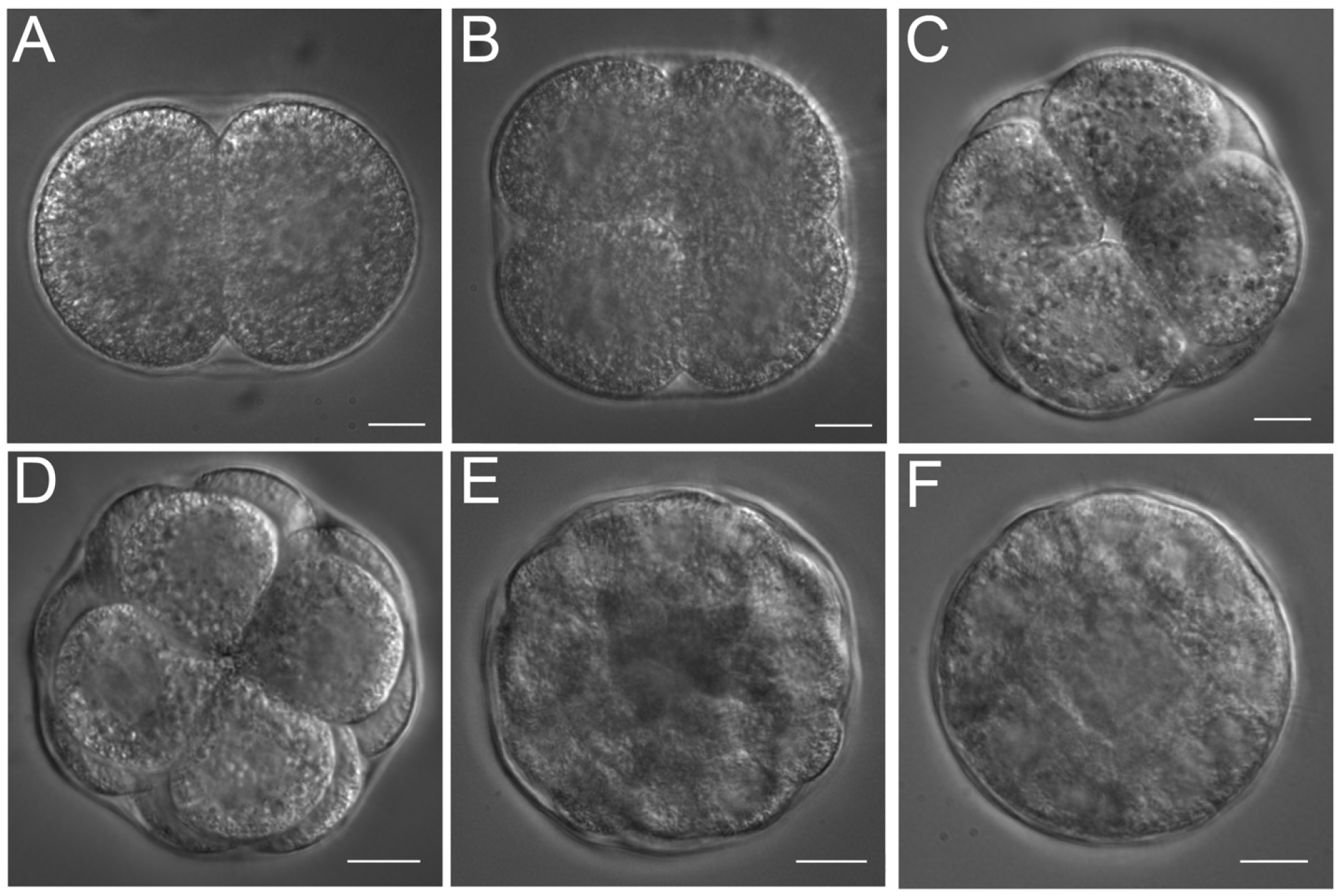
Embryonic cleavage stages in *Hydroides* elegans. All panels are depicted with the animal pole of the embryo pointing out of the page, and the vegetal pole pointing in to the page. Panels A-B and E-F are single equatorial focal planes. Panels C-D are composite images from multiple focal planes. A) 2-cell embryo. B) 4-cell embryo. C) 8-cell embryo. D) 16-cell embryo. The vegetal blastomeres are not in view. E) 32-cell embryo. F) 60-cell embryo. All panels scale=10μm.

The formation of the gut (Figure 4) via invagination at the vegetal pole[16, 47] begins shortly after the completed cleavages of the 64-cell stage embryo, now called a blastula (Figure 4A). The embryos also become ciliated (Figure 4B-C, white arrows) and hatch to begin their free-swimming stages, forming a characteristic apical tuft (Figure 4C, yellow arrow) typical of many marine larvae, along with a thick ciliary band that will eventually form the prototroch[48].

**Figure 4.**
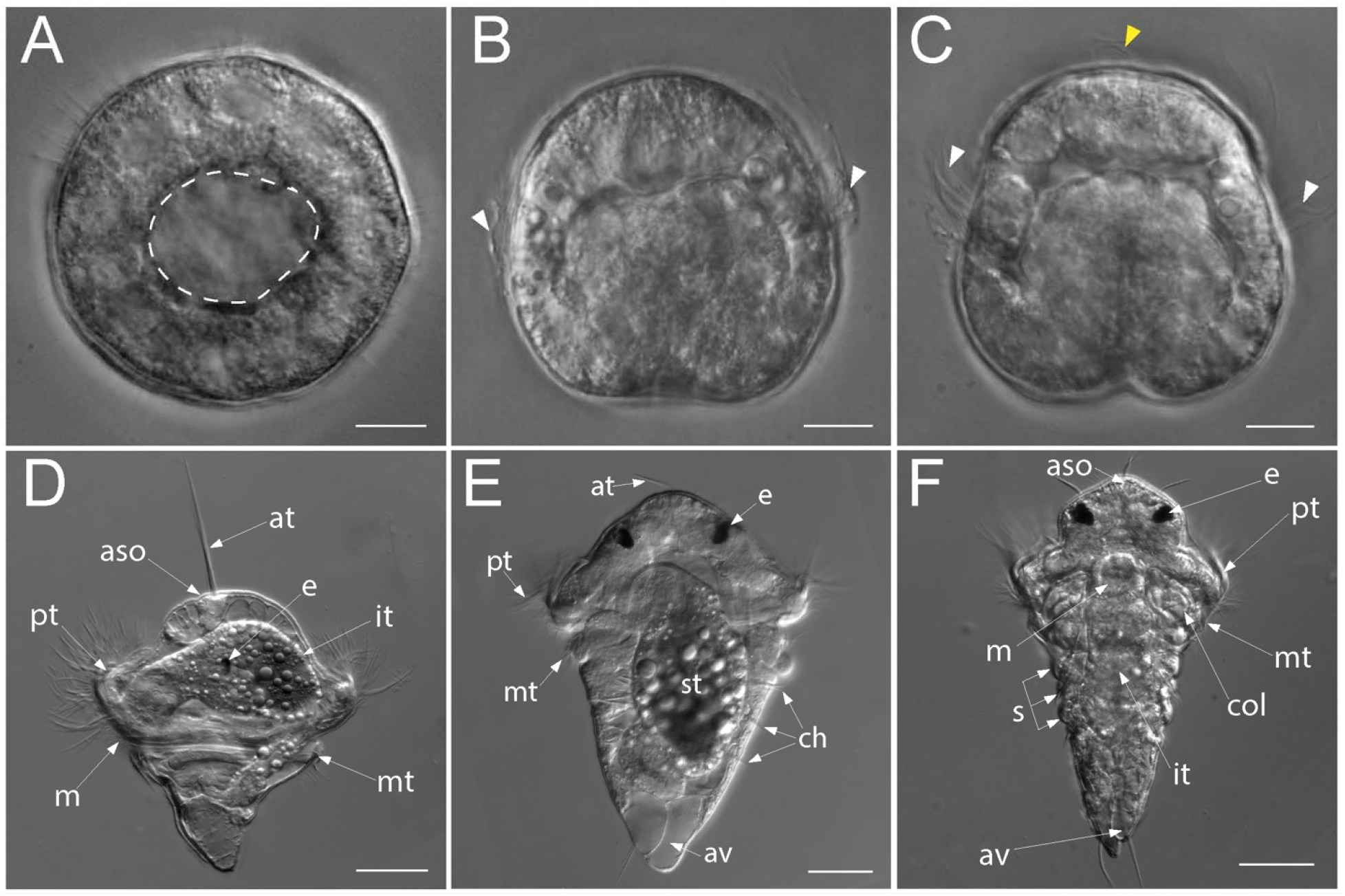
Gastrulation and larval stages in *Hydroides elegans*. A) Ciliated blastula. Dashed white line outlines the blastocoel. B) Mid-gastrula, oral side of the animal is pointing down. White arrows indicate ciliated cells of the developing prototroch. C) Late gastrula, hatched. White arrows mark ciliated cells of the developing prototroch. Yellow arrow indicates cilia of the apical tuft. D) Lateral view of a feeding trochophore larva. E) Aboral view of a segmented nectochaete larva. F) Oral view of a metamorphically competent larva. Labelling for panels D-F: at=apical tuft, aso=apical sensory organ, av=anal vacuole, ch=newly formed chaetae, col=collar, e=eye spot, it=intestinal tract, m=mouth, mt=metatroch, pt=prototroch, s=larval segments with chaetae, st=stomach. For panels A-C scale=10μm, for panels D-F scale=50μm. Panels D-F are from composite images from multiple focal planes.

Prior work has demonstrated that the blastopore partially closes along the ventral midline forming the mouth[16], and the developing gut tube traverses the blastocoel, bending towards the aboral side of the animal[16]. Eventually the epithelium of the developing gut fuses with the aboral ectoderm to form the anus[15, 16, 47, 48]. The complete gut is tripartite, having a differentiated fore-, mid-, and hindgut. Feeding of the trochophore larva (Figure 4D) can then begin at approximately 12 hours post-fertilization (hpf)[17]. The beating of the plush ciliary band forming the prototroch, as well as the opposing accessory band, the metatroch, helps to direct food particles towards the mouth.

Over the course of several days, the trochophore larva feeds. Two eye spots form in sequence. The pigmentation of the first eye spot is visible as early 11 hours post-fertilization, followed by the second which is clearly visible by 72 hpf, and the posterior portion of the body elongates[12]. Three segments arise nearly simultaneously on the posterior portion of the body, each segment develops chaetal sacs and parapodia[12]. The larva is now a segmented nectochaete (Figure 4E), and continues to feed, though no additional segments are added to the body until after metamorphosis. Upon reaching competency (Figure 4F) between 5-6 days post-fertilization, the larva has three complete body segments and looks morphologically distinct from the pre-competent larva. In short, competent larvae are larger in overall body size, and the body profile from the prototroch to the tail is more cone-like in comparison to pre-competent larvae which have a more rhomboid appearance. The head is more clearly defined from the body in competent larvae, because there is not any tapering or gradual sloping at the anterior region of the body, unlike in the pre-competent larvae (Figure 4E). The competent larva is then ready to settle on a substrate and undergo metamorphosis.

Metamorphosis is induced by contact with heterogeneous multispecies or monospecies biofilms[7, 8, 10, 49]. Settlement in response to a natural seawater-derived biofilm is characterized by a slowing of swimming speed, and probing of a surface[17]. While probing the settlement surface, the larva makes transient, reversible contact with the surface and appears to crawl along it, while wiggling or flexing the body by contracting the muscles along the body wall before resuming a slow swimming speed. If suitable, the larva attaches (Figure 5A) itself to a surface and begins to secrete a primary proteinaceous tube[19] (Figure 5B, dashed line) and shed the prototroch cilia (Figure 5C, noted with “*”) which are no longer needed for swimming. Major tissue remodeling occurs as the anterior region of the larva’s body changes shape to form a collar (Figure 5D), food groove cells between the prototroch and metatroch are shed (Figure 5E), and cilia from the metatroch are lost (Figure 5E) as lobes extended laterally (Fig 5F-G) on the head of the animal beginning formation of the branchial crown, and a mineralized secondary tube is deposited.

**Figure 5.**
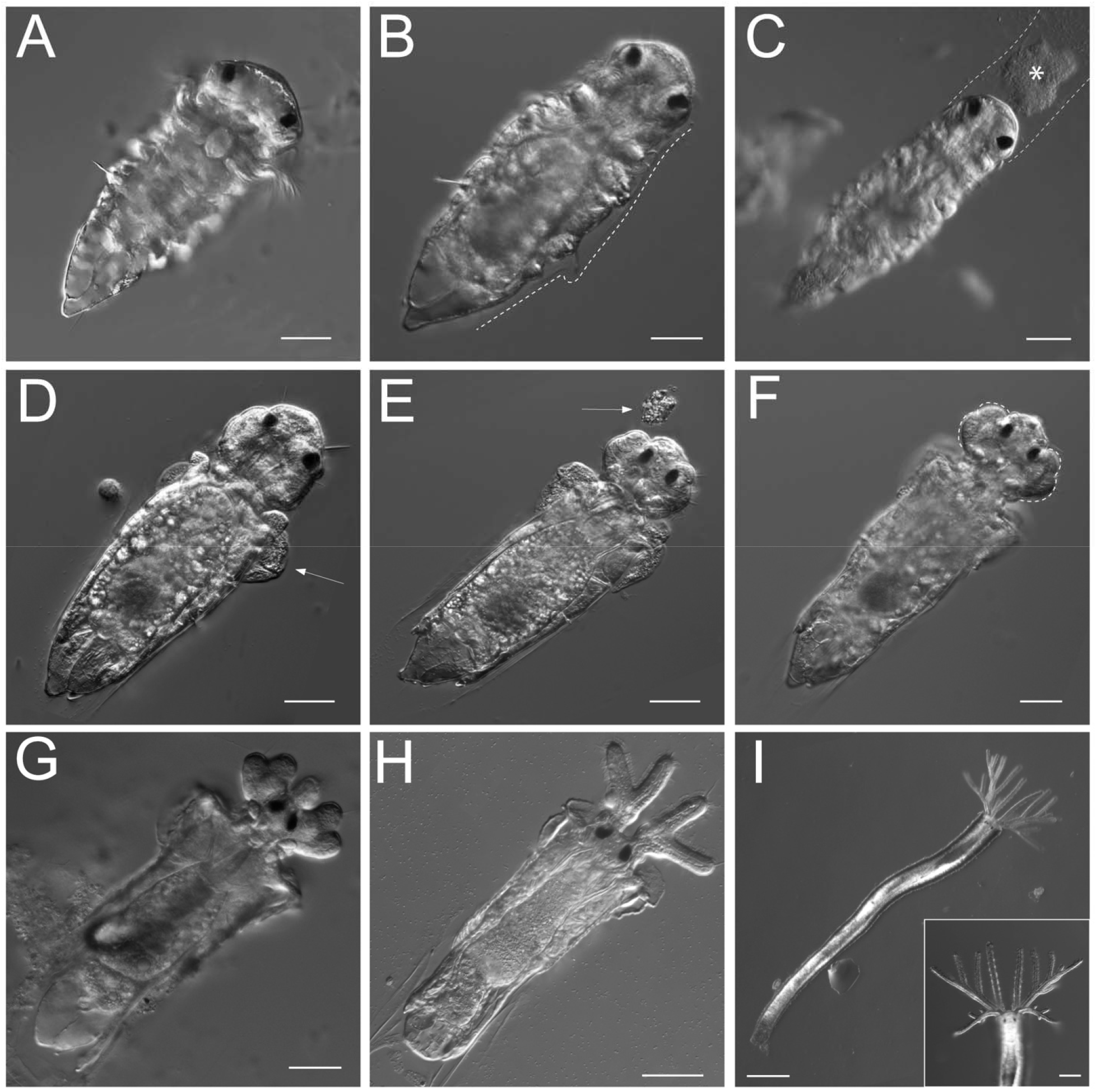
Morphological changes associated with bacteria-stimulated metamorphosis in *Hydroides elegans*. A) Attachment to substrate. B) Formation of the primary tube (indicated by a dashed white line on one side of the larva). C) Shedding of prototroch cilia (*). D) Collar eversion (arrow). E) Shedding of food groove cells (arrow). F) Beginning of lobe formation (dashed white line). G) Lateral extension and elaboration of anterior lobes. H) Juvenile at 24 hours post-metamorphosis. I) Juvenile at 1 week post metamorphosis. Inset of head and elaboration of branchial crown. For panels A-H, scale = 50 μm. For panel I, scale = 250 μm, inset scale = 100 μm.

Post-metamorphic juveniles (Figure 5H-I) continue to elaborate the anterior portion of the body, forming multiple radioles on the branchial crown for feeding (Figure 5I), as well as the operculum in 10-14 days. They continue to deposit their calcareous tubes and within a week of metamorphosis grow to several millimeters in length (Figure 5I, Figure 6). Under ideal culturing conditions with ample food and clean, well oxygenated water, they have a generation time of approximately three to four weeks[11]. In the wild, they settle gregariously, forming dense communities in a matter of weeks, and can be cultured as single tubes in the lab to isolate individual sexually mature adults (Figure 1A).

**Figure 6.**
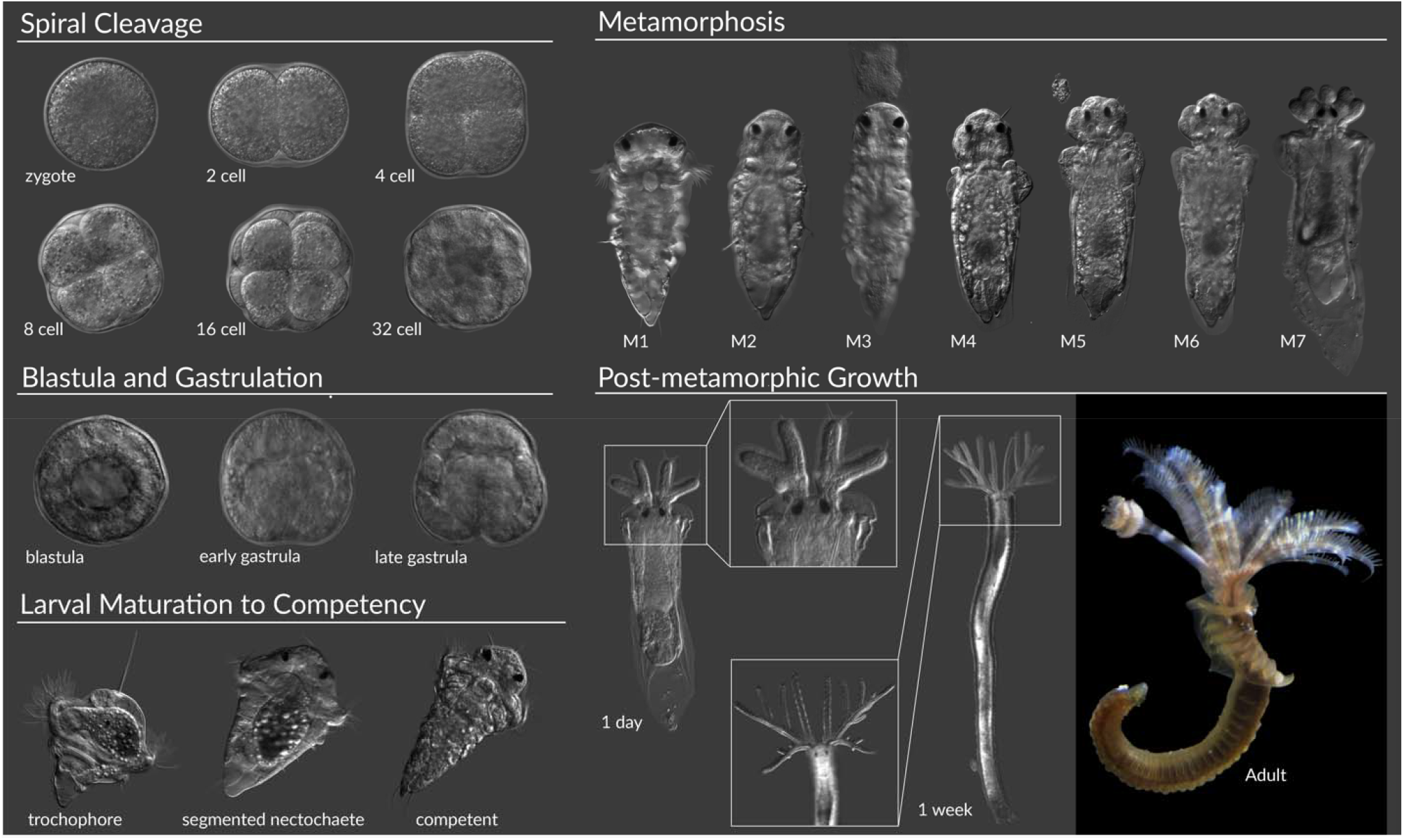
Summary of *Hydroides* development from fertilization through sexual maturation. Major phases of development are labelled with individual stages. Images are not to scale. Metamorphosis stages are labelled M1-M7 and correspond to attachment to substrate (M1), formation of the primary tube (M2), shedding of prototroch cilia (M3), collar eversion (M4), shedding of food groove cells (M5), beginning of lobe formation (M6), lateral extension and elaboration of anterior lobes (M7).

## BACTERIA-STIMULATED METAMORPHOSIS OF HYDROIDES

The role of microbes in mediating animal development is of increasing interest. This is in part due to the diversity of organisms that participate in host-microbe relationships, as well as the diversity of developmental outcomes that arise from interactions with bacteria. The transition to multicellularity[1] as well as switches between asexual to sexual reproductive strategies[2] in choanoflagellates, maturation of the gut in zebrafish[3] and mammals[4], organogenesis in the Hawaiian bobtail squid[5], and initiation of metamorphosis in marine invertebrates[6, 7] including *Hydroides*[8, 9] are just some of the varied ways in which microbial interactions influence eukaryotic development.

Given the remarkable impact that marine bacteria have on the life cycle of *Hydroides*, it comes as no surprise that much attention has been given to characterizing the nature of this host-microbe association during metamorphosis. Many different single strains of bacteria induce *Hydroides* metamorphosis[8]. For example, the Gammaproteobacterium, *Pseudoalteromonas luteoviolacea*, and the Bacteroidetes, *Cellulophaga lytica* [50]. The diverse inductive bacteria also produce diverse bacterial products that promote the same developmental endpoints[51]. For example, *P. luteoviolacea* produces a contractile injection system[52] that is related to the contractile tails of bacterial viruses (bacteriophage). These syringe-like structures are assembled into arrays and deliver a protein payload[53], the effector Mif1, to *Hydroides* larvae which then interacts with host-derived signaling networks[54] to induce metamorphosis. Other bacteria stimulate metamorphosis through different mechanisms[51]. For example, lipopolysaccharide from outer membrane vesicles of some bacteria induce *Hydroides* metamorphosis[50]. These biochemically unique products that induce metamorphosis from diverse bacteria open up a future for discovery. The molecular mechanisms driving metamorphosis in *Hydroides* are likely to reveal fundamental means of interaction at play between hosts and microbes across the tree of life.

Working in the *Hydroides* model system raises interesting questions about the nature of this marine tubeworm-microbe relationship: 1) do other diverse bacterial cues for metamorphosis exist?; 2) given the heterogeneity of marine biofilms, what are the respective contributions of individual bacteria within it to metamorphosis, effectively how potent are the various cues, are there combinatorial effects of these products, and are they distinct enough to be differentiated by the animal?; and 3) do these diverse bacterial cues actually yield identical developmental outcomes, or does the mechanism of metamorphosis initiation influence the sequence of signaling and morphogenetic events[17], and if so, how?

## THE FUTURE OF HYDROIDES

As the *Hydroides* model continues to gain traction in the research community, it is clear there are many areas with potential for discovery. This includes comparative evolutionary and developmental biology, genetic tool development, biotechnology, and the discovery of bacteria-derived natural products.

*Hydroides* can offer unique perspectives on the establishment of host-microbe interactions. Developmental regulatory networks have yet to be integrated with components from microbial symbionts. In fact, the animal sensing machinery responsible for detecting and responding to bacteria by initiating metamorphosis is not well-resolved for *Hydroides*, or any other animal aside from the highly conserved signaling systems (e.g., PKC and MAPK[9, 54-57]). Thus, there exists a conspicuous gap in knowledge that this Annelid model is primed to fill.

Comparative approaches also allow for the investigation of overlap or repurposing of molecular machinery that drives metamorphosis and facilitates host-microbe interactions. For example, immune genes have rapidly evolved and function primarily to detect and respond to encounters with bacteria. However, repurposing of these components could facilitate interactions with non-pathogenic bacteria and intersect with developmental pathways like those activated during metamorphosis[9, 58]. The *Hydroides* genome contains more genes in common with humans than *C. elegans* does[9], placing *Hydroides* as a model system with the potential for discoveries relating to human development, health, and disease.

The *Hydroides* model sits squarely at the junction of questions in ecology, evolution, and development. As a broadcast spawning marine invertebrate that has a vast dispersal range, we can gain insight into how animal ecology is influenced by environmental microbes that stimulate metamorphosis. This would be applicable, for example, in reef environments where recruitment and successful settlement of dispersed pelagic larvae is essential for habitat maintenance. These eco-evo-devo perspectives[59] may illuminate the evolution of host-microbe symbioses, and also add a new layer of context to ecosystem dynamics and management for vulnerable marine habitats.

A clear and current bottleneck in the development of *Hydroides* as a model system for host-microbe interactions is a lack of established techniques for genetically manipulating the *Hydroides* host. There has been recent success in the genetic manipulation of related Annelids such as *Platynereis dumerilii*[60, 61] and *Capitella teleta*[62, 63] and we are optimistic that *Hydroides* will be amenable to genetic manipulation. Improvements in husbandry and long-term culture will facilitate generation of animal lines in *Hydroides*, enabling *Hydroides* to keep pace with advancements in other marine invertebrate larvae[64-69]. In parallel, emerging synthetic biology tools in established model microbes also hold promise for the manipulation of more diverse marine bacteria that promote *Hydroides* development[70]. These new tools compliment powerful classical bacterial genetics approaches which have already helped reveal key aspects of *Hydroides* interaction with stimulatory microbes[52, 53].

Future work leveraging *Hydroides* holds promise for promoting biotechnology in the contexts of evolution, the environment, and human health. For example, insights gained from *Hydroides* to understand bacteria-stimulated metamorphosis could aid in the husbandry of invertebrates in aquaculture[71]. *Hydroides* has been studied intensively as a biofouling organism[11] and research on antifouling technologies could provide significant economic relief for shipping and naval sectors[43]. The products bacteria produce to promote *Hydroides* metamorphosis and the machinery *Hydroides* uses to sense these products could lead to broader discoveries about mechanisms and tools for host-microbe interactions, and how these interactions evolve on a genetic and chemical level. For example, we found that gene clusters encoding Contractile Injection Systems originally discovered to promote *Hydroides* metamorphosis are strikingly similar to gene clusters found in *Bacteroidales* bacteria from healthy human microbiomes[72]. Moreover, understanding the payload delivery and targeting mechanism of the MACs produced by *P. luteoviolacea* could enable new types of therapeutic delivery systems[73].

## CONCLUSION

Environmental bacteria are becoming increasingly recognized for their significant roles in regulating health and development[6, 74]. The utilization of the *Hydroides*–microbe model offers opportunities in the eco-evo-devo landscape to understand the conservation and diversity of microbial drivers of eukaryotic development and disease states.

## EXPERIMENTAL PROCEDURES

### Husbandry, Spawning and Fertilization

Adult worms are collected from Quivira Basin in Mission Bay, San Diego CA. Animals are housed in a suspended basket in a recirculating tank at room temperature, supplied with artificial filtered sea water (AFSW) at 35 PSU. Adults are fed *ad libidum* with a monoculture of the marine algae *Isochrysis galbana* (Carolina Biological: Item #153180). To spawn animals, individual adult tubes are selected and placed in separate glass petri dishes (60mm x 15mm) filled with AFSW. Using forceps, the tubes are cracked, revealing the animal inside which releases thousands of eggs or streams of sperm upon disturbance.

Eggs are collected with a trimmed plastic pipette tip and washed into fresh FASW to minimize debris left over from tube fracture. Sperm is collected with a pipette in minimal sea water, and then diluted further into a fresh sperm suspension for fertilization. Eggs are diluted to ∼500/mL and 4-5 drops of sperm suspension is added. The dish is swirled to mix. Ten minutes after the addition of sperm to the eggs, the eggs are collected and washed into fresh FASW three times to eliminate excess sperm and minimize the risk of polyspermy. Eggs and embryos are monitored over the next several hours to ensure fertilization since no obvious morphological changes occur at fertilization. All cultures of embryos are maintained at a density of 1 larva/ml in an incubator at 25°C, with foil covers to prevent evaporation. Larval cultures have water changes daily starting at Day 2, and are fed *I. galbana* daily at a density of 6×10^4^ cells/mL. At 5 days post-fertilization, the majority of larvae have reached competency (there is some asynchrony due to variation in individual larval feeding efficiency) and are transferred to a dish that has a natural biofilm coating the bottom of the dish. The natural biofilm is generated by filling the dish with raw seawater from the worm’s natural habitat and allowing that to sit for four to five days. The worms metamorphose over the course of ∼12 hours upon interacting with the natural biofilm, and are then either imaged or kept for longer-term culture to adulthood. Metamorphosed worms that are kept for culture to adulthood are fed *I. galbana* every other day *ad libidum*, and water changes are completed on an identical schedule.

### Live-cell Imaging of Development

All stages except the adult of *Hydroides elegans* were imaged live on a Zeiss Axio Observer.Z1 using objectives ranging from 5x to 100x oil immersion, all with differential interference contrast (DIC) optics. Images were captured using the ZEN software suite, and measurements were added using Fiji (National Institutes of Health, Bethesda, Maryland)^16^. Composite DIC images were rendered from Z-stacks of animals using Helicon Focus Pro Unlimited (v6.8.0, Helicon Soft Ltd.). Adult animals were imaged live on a Wild Heerbrugg M5a stereo microscope fitted with a QImaging R3 Retiga CCD camera.

## Acknowledgements

We would like to thank the members of the Shikuma Lab for constructive feedback on the present manuscript. This work was supported by the National Science Foundation (1942251), the Gordon and Betty Moore Foundation (GBMF9344; https://doi.org/10.37807/GBMF9344), Office of Naval Research (N00014-20-1-2120), the National Institutes of Health, NIGMS (R35GM146722) and the Alfred P. Sloan Foundation, Sloan Research Fellowship.

